# The white matter fiber tract deforms most in the perpendicular direction during *in vivo* volunteer impacts

**DOI:** 10.1101/2024.03.26.585293

**Authors:** Zhou Zhou, Christoffer Olsson, T. Christian Gasser, Xiaogai Li, Svein Kleiven

**Affiliations:** Division of Neuronic Engineering, KTH Royal Institute of Technology, Stockholm, 14152, Sweden; Division of Biomedical Imaging, KTH Royal Institute of Technology, Stockholm, 14152, Sweden; Material and Structural Mechanics, Department of Engineering Mechanics, KTH Royal Institute of Technology, Stockholm, 14152, Sweden

**Keywords:** Volunteer head impacts, tagged magnetic resonance imaging, diffusion tensor imaging, tract-related strains, *in vivo* white matter fiber deformation

## Abstract

White matter (WM) tract-related strains are increasingly used to quantify brain mechanical responses, but their dynamics in live human brains during *in vivo* impact conditions remain largely unknown. Existing research primarily looked into the normal strain along the WM fiber tracts (i.e., tract-oriented normal strain), but it is rarely the case that the fiber tract only endures tract-oriented normal strain during impacts. In this study, we aim to extend the *in vivo* measurement of WM fiber deformation by quantifying the normal strain perpendicular to the fiber tract (i.e., tract-perpendicular normal strain) and the shear strain along and perpendicular to the fiber tract (i.e., tract-oriented shear strain and tract-perpendicular shear strain, respectively). To achieve this, we combine the three-dimensional strain tensor from the tagged magnetic resonance imaging (tMRI) with the diffusion tensor imaging (DTI) from an open-access dataset, including 44 volunteer impacts under two head loading modes, i.e., neck rotations (N = 30) and neck extensions (N = 14). The strain tensor is rotated to the coordinate system with one axis aligned with DTI-revealed fiber orientation and then four tract-related strain measures are calculated. The results show that tract-perpendicular normal strain peaks are the largest among the four strain types (*p* < 0.05, Friedman’s test). The distribution of tract-related strains is affected by the head loading mode, of which laterally symmetric patterns with respect to the midsagittal plane are noted under neck extensions, but not under neck rotations. Our study presents a comprehensive *in vivo* strain quantification towards a multifaceted understanding of WM dynamics. We find the WM fiber tract deforms most in the perpendicular direction, illuminating new fundamentals of brain mechanics. The reported strain images can be used to evaluate the fidelity of computational head models, especially those intended to predict fiber deformation under non-injurious conditions.

## Introduction

Traumatic brain injury (TBI) is a pressing public health and socio-economic problem. Annually, around 69 million people worldwide suffer a TBI.^1^ In Europe, about 7.7 million people live with TBI-induced disability, incurring immense personal suffering and huge economic costs.^2^ In Sweden, the annual number of TBI-related emergency department visits is estimated to be 30,000 with 10,000 requiring hospitalization.^3^ Despite substantial efforts to reduce the occurrence and mitigate the severity, TBI remains to be the leading cause of death in young adults and its incidence is alarmingly increasing in the elderly.^4^ There is a clear need for more research to address this TBI-related epidemic, of which how the brain responds during impacts is the central question to be answered.

Various approaches are used to elucidate the biomechanical fundamentals behind brain trauma. In *in situ* cadaveric tests, the post-mortem human subjects are meticulously instrumented to measure brain responses with the impact severity close to the traumatic level.^5^ For example, Hardy and colleagues^6, 7^ conducted 35 cadaveric tests with blunt impacts to track the motion of 12 sparse makers within the deep brain region, which was later analysed to estimate the brain strain^8, 9^ and strain rate.^10^ More recently, Alshareef and coworkers^11^ instrumented six head-neck complexes, each of which equipped 24 markers throughout the brain parenchyma, to obtain brain motion in 72 impacts with pure rotational loadings. *In vitro* models play a crucial role in comprehending the mechanobiological cascades of TBI, especially at the cellular and subcellular level, by relating controlled mechanical inputs to the resultant morphological damage or electrical dysfunction of experimental tissues.^12, 13^ For example, Morrison III and colleagues mechanically injured *in vitro* cultured rat hippocampi. It was found that the cell death was monotonically dependent on the strain,^14^ while the functional tolerance was affected by strain and strain rate in a complex fashion.^15^ *In silico* head models are increasingly used to quantitatively simulate the localized tissue response during head impacts.^16–18^ Modern *in silico* models feature subject-specific geometry,^19, 20^ anisotropic and heterogeneous brain material,^21–23^ and fluid representation of the cerebrospinal fluid.^24–26^ Encouraging correlations have been reported between the response of the *in silico* model and experimentally quantified histology or clinically diagnosed injury.^27–30^

Another important line to study human brain biomechanics is *in vivo* quantitative imaging, which can offer complementary information of the intracranial dynamics under non-injurious conditions.^31^ Today, many *in vivo* imaging modalities (e.g., magnetic resonance elastography (MRE),^32, 33^ amplified magnetic resonance imaging (MRI),^34^ and phase-contrast MRI)^35^ are used to enlighten the brain mechanics under volunteer impacts or physiological conditions, of which the tagged MRI (tMRI) is particularly illuminating. tMRI is an imaging technique that was originally developed to measure cardiovascular deformation.^36^ In 2005, Bayly and colleagues^37^ pioneeringly implemented tMRI to image brain deformation under sub-injurious occipital impacts. Over the past two decades, tMRI has been applied to measure brain responses in different voluntary loading scenarios (e.g., axial rotation,^38^ frontal impact)^39^ with multiple technical advancements to improve imaging efficacy, accuracy, and resolution.^40–43^ In 2021, Bayly and colleagues^44^ established an open-access dataset, i.e., the Brain Biomechanics Imaging Resource (BBIR) dataset that includes tMRI-recorded brain responses from 50 volunteer tests and other pertinent information (e.g., impact kinematics, structure imaging of volunteers).

Axonal injury is a frequent form of trauma in the white matter (WM) and characterized by diffuse lesions within the fiber tracts usually consisting of oriented axons.^45, 46^ Given this pathologic feature, there has been growing interest in quantifying the loading endured by the fiber tract.^47^ For *in vivo* experimental data, only one study^48^ based on the BBIR dataset calculated the normal strain along the fiber tracts (i.e., tract-oriented normal strain, alternatively termed as fiber strain,^49^ tract-oriented strain,^50–52^ and axonal strain^49, 53^ in the literature) by projecting the tMRI-recorded brain strain tensor along the primary fiber direction estimated from the diffusion tensor imaging (DTI) in 20 volunteer impacts (10 neck rotations and 10 neck extensions). DTI is an imaging technique in which the anisotropic movement of water molecules within the brain is utilized to estimate the WM fiber tract.^54^ Despite interesting results, the tract-oriented normal strain only represents one aspect of fiber deformation. Considering the tensorial nature of the strain and the complexity of impact loading, it is rarely the case that the fiber only endures the tract-oriented normal strain. A multifaceted quantification of fiber deformation remains to be performed.

The current study aims to present a comprehensive quantification of WM tract-related deformation under volunteer impacts. To this end, the tMRI-recorded brain strain tensor data in 44 volunteer tests is related to the DTI-derived fiber direction. These volunteer tests and relevant data are acquired from the open-accessed BBIR dataset.^44, 48^ By combining multimodal imaging analyses with continuum mechanics, we quantify the normal strain and shear strain along and perpendicular to the fiber tract. This investigation finds the WM tract deforms most in the perpendicular direction, unravelling new fundamentals of brain mechanics. The reported strain data can be used to evaluate the biofidelity of *in silico* models.

## Methods

### Volunteer tests and imaging data

The brain strain tensor data during volunteer tests and brain imaging are acquired from the publicly available BBIR dataset where details regarding study approval, subject recruitment, test protocol, imaging acquisition, and processing are available elsewhere.^44^ Briefly, the volunteer was rapidly rotated at non-injurious levels by a custom-built neck rotation device (rotation within the axial plane along the inferior-superior axis) or neck extension device (rotation within the sagittal plane about the left-right axis). The angular kinematics within the primary rotating plane were recorded, i.e., the axial plane for the neck rotation (Fig 1A) and the sagittal plane for the neck extension (Fig 1B). In each test, three-dimensional (3D), full-field, time-varying brain strain tensor, including six independent components (Fig 1C), were measured using tMRI at a spatial resolution of 1.5 mm or 2.0 mm and a temporal resolution of 18 ms, 19.5 ms or 20 ms. These subject-specific tensorial brain strain data under known head loading conditions were further complemented by multimodal structural imaging data of the same subject. As the WM is the structure of interest in the current study, DTI and brain segmentation are particularly relevant. In the dataset, DTI-derived information, including the principal eigenvector of the diffusion tensor and the fractional anisotropy (FA) map (Fig 1D), were available. The brain was segmented into different anatomical structures (Fig 1E), of which the WM region included cerebral WM, cerebellar WM, and brainstem. For each test, the structural images were processed with the same spatial resolutions as the tMRI.

**Fig 1.**
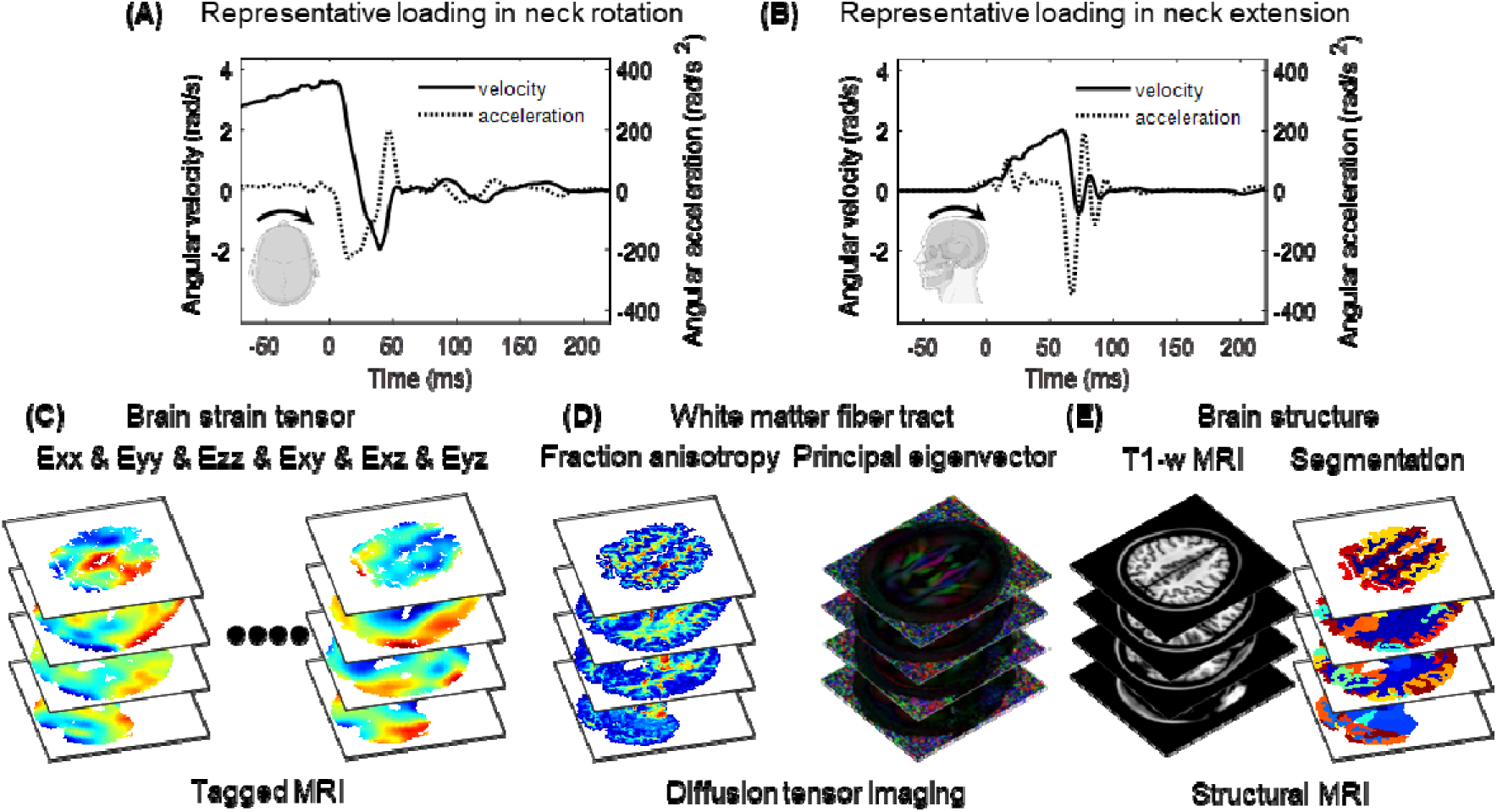
Data from voluntary tests in neck rotation and neck extension and imaging acquisition obtained from the Brain Biomechanics Imaging Resource dataset (https://www.nitrc.org/projects/bbir/).^44, 48^ (A) Angular kinematics in the axial plane for one representative neck rotation test. (B) Angular kinematics in the sagittal plane for one representative neck extension test. (C) 3D full-field, time-varying Green-Lagrange brain strain tensor measured by tagged MRI. (D) White matter fiber-related information (i.e., fractional anisotropy, and principal eigenvector quantifying the primary direction of fiber tracts) derived from diffusion tensor imaging. (E) Brain structure and segmentation. Note that the kinematics in subfigures A and B correspond to the two impacts that are used as representative tests for illustration purposes in the “Results” section.

In the current study, 44 volunteer tests are included to estimate the WM tract-related strains. Note that, from the 50 tests in the original dataset, 4 tests performed on cadavers and 2 tests without DTI are excluded. These 44 tests, including 30 neck rotations and 14 neck extensions, were acquired from 38 subjects (20 male, 18 female) with an average age of 31.7 ± 10.8 years (range: 19-59 years). Structural imaging data (e.g., brain segmentation, DTI, and its derivatives) are spatially aligned with the tMRI-derived brain strain tensor data, except for 4 tests with imaging misalignment. Co-registrations between different imaging modalities are performed on these 4 tests, with the use of common neuroimaging tools from the FMRIB Software Library.^55, 56^

### Computation of white matter tract-related strains

To calculate the WM tract-related deformation, the tMRI-derived brain strain tensor is rotated to the coordinate system with one axis aligned with the DTI-derived fiber orientation (referred to as tract-aligned coordinate system), through which the components in the transformed tensor quantify multiple strain modes endured by the fiber (Fig 2). This calculation is applied to all WM voxels (i.e., voxels with FA values ≥ 0.2), of which the fiber orientation is approximated as the first principal eigenvector of the diffusion tensor. In each time frame of tMRI, we calculate the tract-oriented normal strain ( ), a metric that has also been chosen as the parameter of interest in one early *in vivo* study.^48^ As a step further, we also study three new tract-related strains, including tract-perpendicular normal strain ( , the maximal normal strain perpendicular to the fiber tract), tract-oriented shear strain ( , the shear strain along the fiber tract), and tract-perpendicular shear strain ( , the maximal shear strain perpendicular to the fiber tract). As magnitudes of strain components are dependent on the choice of tract-aligned coordinate systems, two notations, and , are introduced, representing their maximum values. This calculation is performed at each time frame of the tMRI. The mathematical derivation of tract-related strains is detailed in Appendix A.

**Fig 2.**
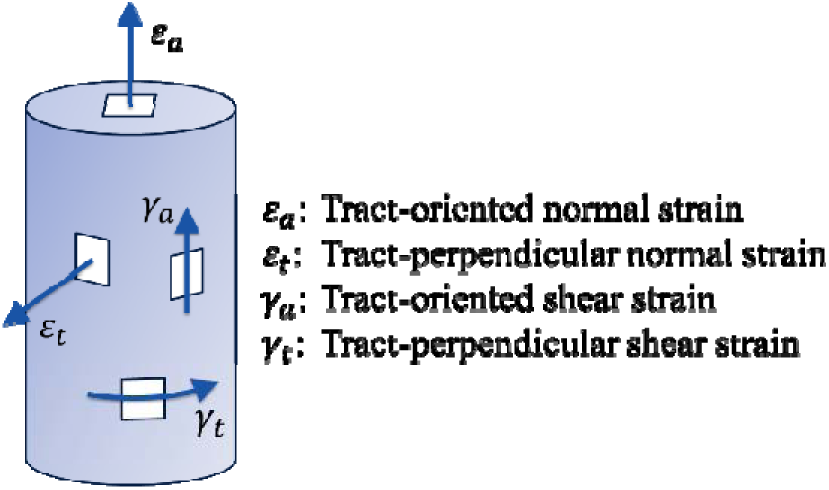
Sketch of four tract-related strains characterizing the normal strain and shear strain along and perpendicular to the fiber tract.

For the four tract-related strains, we identify the peak values accumulated over time during the two head loading protocols (i.e., neck rotation and neck extension). This is motivated since the time-accumulated strain peaks are often used for injury analysis.^57, 58^ It is also a compromised strategy to address the disparity of temporal resolution in tMRI (30 tests with a time resolution of 18 ms vs. 3 tests with a time resolution of 19.5 ms vs. 11 tests with a time resolution of 20 ms). For the spatial resolution disparity of tMRI (31 tests with a voxel resolution of 1.5 mm vs. 13 tests with a voxel resolution of 2 mm), its influence on the tract-related strains remains negligible, see Appendix B. Hence, strain results with different temporal and spatial resolutions are combined for the statistical analysis. For each test, the peak tract-related strains are computed for three WM subregions (i.e., cerebral WM, cerebellar WM, brainstem) and the whole WM. To avoid potential artifacts, the 95^th^ percentile peak values are reported, the same strategy as in previous studies.^48, 59^

### Statistical analysis

Whether magnitudes of four tract-related strains (non-normally distributed) differ are tested using the Friedman’s test with the Bonferroni correction for multiple comparisons to determine if differences between individual strain pairs are statistically significant. This test is performed at the regional level (i.e., cerebral WM, cerebellar WM, brainstem) and the whole WM level. To compare the fiber deformation between two head loading modes (i.e., neck rotation and neck extension), Mann-Whitney U tests are conducted on four tract-related strain peaks. Within each head loading mode, we perform Pearson’s correlations to test the dependency of four tract-related strains at 4 regions of interest (i.e., cerebral WM, cerebellar WM, brainstem, and whole WM) on two angular kinematic peaks (i.e., peak change in angular velocity and peak angular acceleration). 32 Pearson’s correlations are conducted for a given head loading mode with the Benjamini-Hochberg method implemented to consider the correction for multiple comparisons.

## Results

At the individual level, tract-related strain responses in one specific subject, who performed both a neck rotation and a neck extension, are reported in Fig 3 as illustrative cases. The peak change in angular velocity and peak angular acceleration are 5.6 rad/s and 226.8 rad/s^2^ for the neck rotation (Fig 1A), and 2.8 rad/s and 345.6 rad/s^2^ for the neck extension (Fig 1B). Spatially heterogeneous strain patterns are noted in the WM and the regions with high strains (i.e., areas in red in Fig 3A and Fig 3C) are dependent on the head loading mode and strain type. For neck rotation, high tract-oriented normal and shear strains are sparsely distributed and mostly occur at the boundary of cerebral WM and periventricular region, while high tract-perpendicular normal and shear strains exhibit relatively continuous patterns that extend from the anterior-superior site in the left cerebrum to the deep region in the right cerebrum (Fig 3A). In the neck extension, high tract-oriented normal strain is primarily noted in the brainstem, while high tract-perpendicular normal and shear strains, tract-oriented shear strain appear isolated and occur mostly in the anterior-superior cerebral region (primarily close to the midsagittal plane), inferior cerebellar area, and brainstem (Fig 3C). For the strain peak in the neck rotation (Fig 3B) and neck extension (Fig 3D), the largest value is consistently noted in tract-perpendicular normal strain across all examined regions (i.e., cerebral WM, cerebellar WM, brainstem, and whole WM).

**Fig 3.**
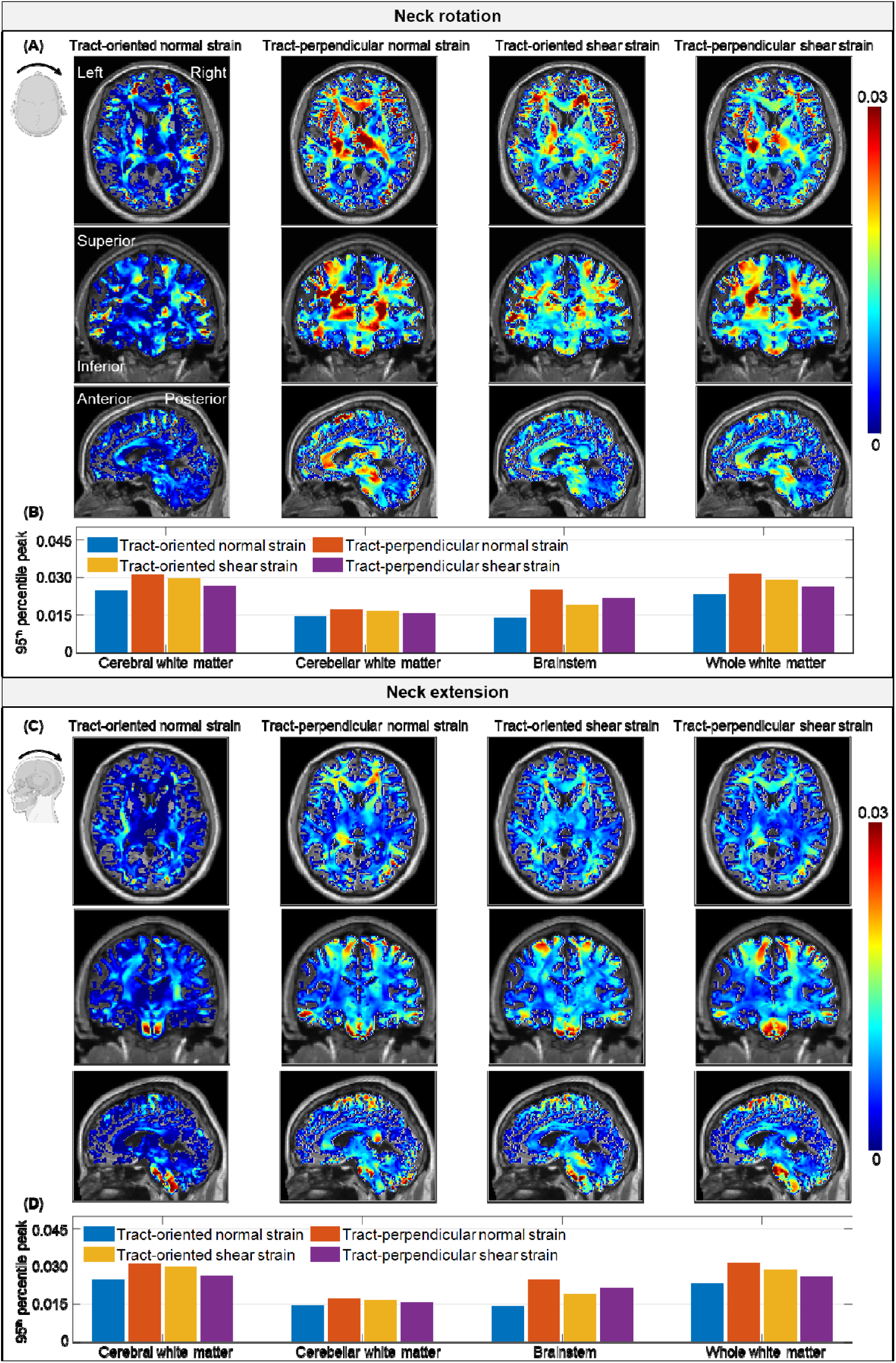
Time-accumulated tract-related strain peaks in one neck rotation (A-B) and one neck extension (C-D) that were conducted on the same subject. (A) Axial, coronal, and sagittal views of tract-oriented normal strain, tract-perpendicular normal strain, tract-oriented shear strain, tract-perpendicular shear strain (color), and T1-w MRI (grayscale) during neck rotation with 95^th^ percentile peak values in three white matter subregions and the whole white matter shown in subfigure B. Similar plots are presented for neck extension in subfigures C-D. The anatomical orientation is specified in the tract-oriented normal strain contour during neck rotation in subfigure A.

Left-right symmetric strain patterns with respect to the midsagittal plane are noted in the neck extension (Fig 3C), but not in the neck rotation (Fig 3A). This is further quantified by calculating the strain peaks in the left and right brain hemispheres, respectively (Fig 4). In the neck rotation, the strain differences (in absolute percentage) between the left and right brain hemispheres ranges from 2.7% to 34.8%, while in the neck extension, the percentage of inter-hemispheric strain differences varies from 0.3% to 15.4%.

**Fig 4.**
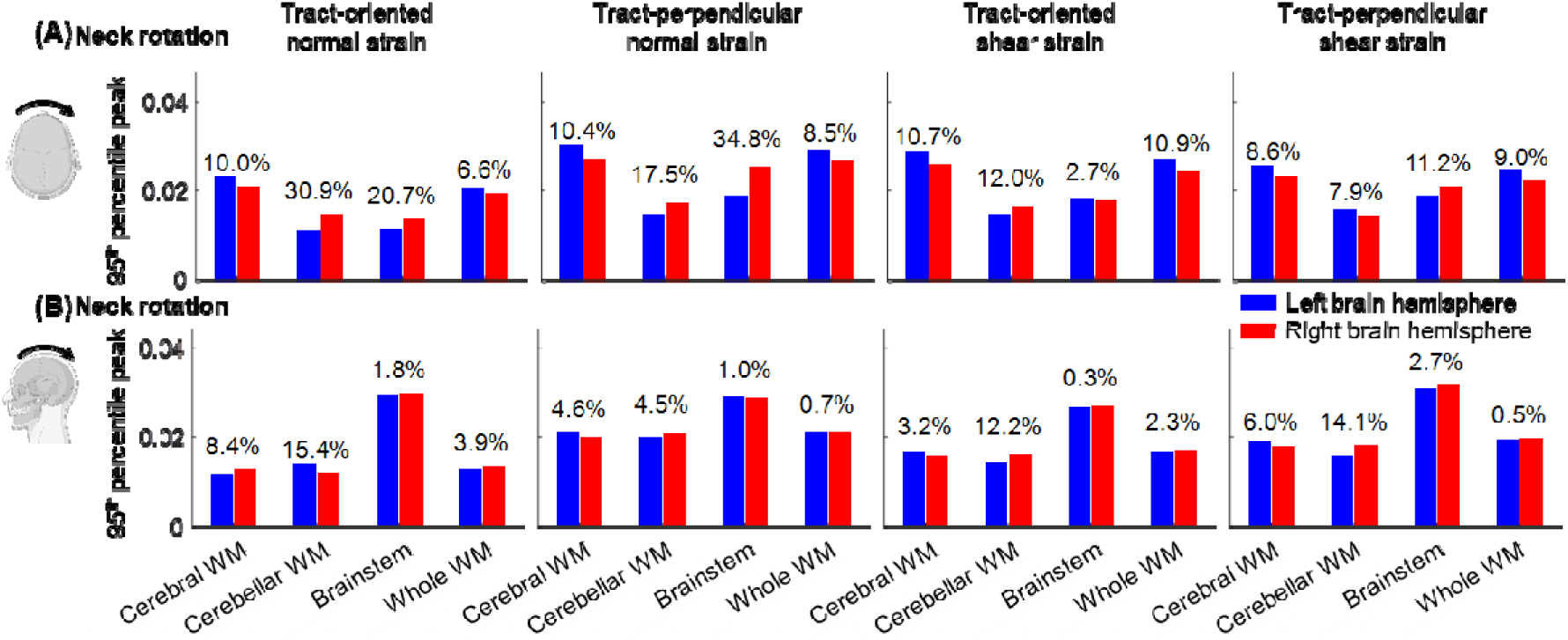
Comparisons of time-accumulated tract-related strain peaks between the left and right brain hemispheres in one neck rotation (A) and one neck extension (B) at three white matter (WM) subregions (i.e., cerebral WM, cerebellar WM, brainstem) and the whole WM level. For each comparison, the strain difference in absolute percentage is labelled with the result from the right brain as the baseline.

At the group level, comparisons among four tract-related strain peaks are summarized in Fig 5. Across all regions of interest, significant differences are noted in magnitudes of tract-related strains (*p*<0.05, Friedman’s tests with the Bonferroni correction of multiple comparisons) with the tract-perpendicular normal strain peak being the largest. The tract-oriented normal strain is significantly smaller than the tract-oriented shear strain and tract-perpendicular shear strain in the cerebral WM, cerebellum WM, and whole WM level, but not in the brainstem. No significant differences are noted between the tract-oriented shear strain and tract-perpendicular shear strain in all examined regions. Detailed plots of inter-strain comparison are presented in Appendix C.

**Fig 5.**
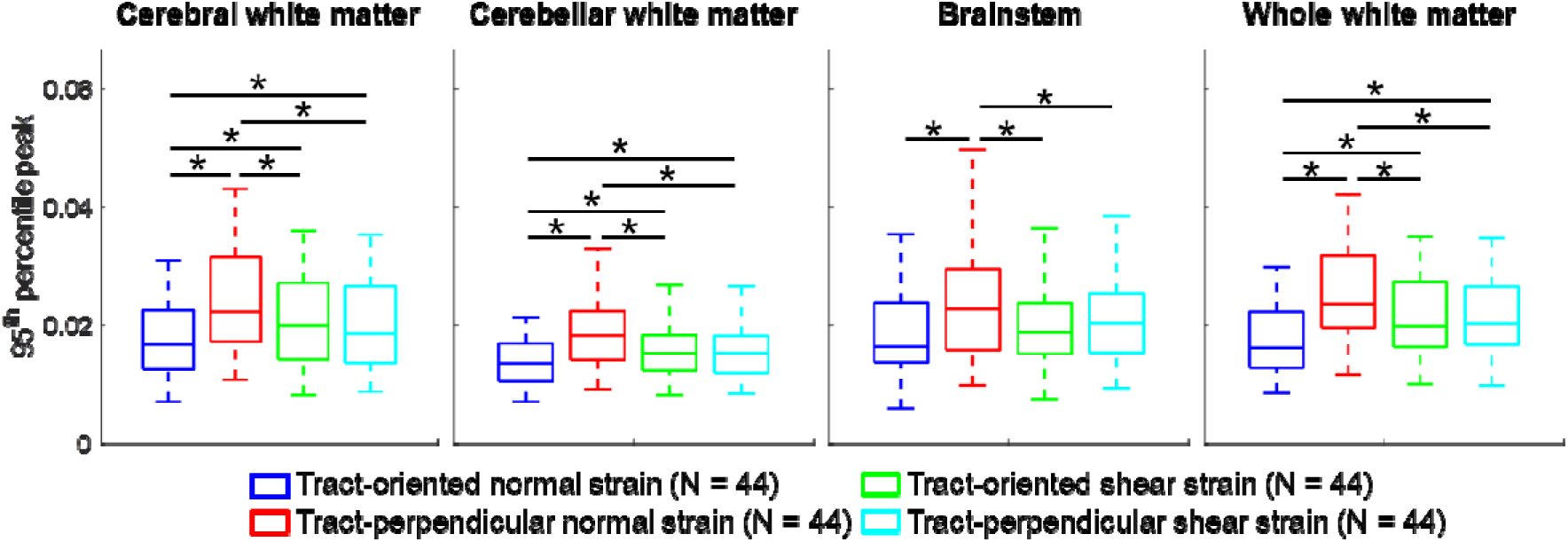
Comparisons of four tract-related strain peaks (95^th^ percentile values) in cerebral white matter (WM), cerebellar WM, brainstem, and the whole WM (* indicates the difference in the referred strain pair is statistically significant when assessed by the Friedman’s tests with the Bonferroni correction of multiple comparisons).

Comparisons of tract-related strain peaks between neck rotations and neck extensions is analyzed in Fig 6. The average values for the peak change in angular velocity and peaking angular acceleration are 4.2 ± 1.2 rad/s (range: 1.8-6.4 rad/s) and 185.5 ± 59.2 rad/s^2^ (range: 49.8-317.0 rad/s^2^) for neck rotations, and 2.1 ± 0.5 rad/s (range: 1.0∼2.8 rad/s) and 228.6 ± 8.3 rad/s^2^ (range: 63.8-345.6 rad/s^2^) for neck extensions. A significant difference (*p*<0.001) is noted for the peak change in angular velocity between two head loading modes (Fig 6A). A summary of tract-related strains in different WM areas is reported in Fig 6B. At the whole WM level, 95^th^ percentile maximum tract-related strains (mean ± standard deviation (range)) in neck rotations are 0.020 ± 0.005 (0.009-0.030) for tract-oriented normal strain, 0.028 ± 0.007 (0.011-0.042) for tract-perpendicular normal strain, 0.024 ± 0.006 (0.010-0.035) for tract-oriented shear strain, and 0.023 ± 0.006 (0.010∼0.035) for tract-perpendicular shear strain. For neck extensions, the maximum tract-oriented normal, tract-perpendicular normal, tract-oriented shear, and tract-perpendicular shear strains are 0.013 ± 0.002 (0.009-0.017), 0.019 ± 0.004 (0.012-0.024), 0.016 ± 0.003 (0.010-0.020), and 0.017 ± 0.003 (0.011-0.022), respectively. When comparing the strain peaks between neck rotations and neck extensions, significant differences (*p*<0.001) are noted in the cerebral WM and whole WM, independent of the strain type.

**Fig 6.**
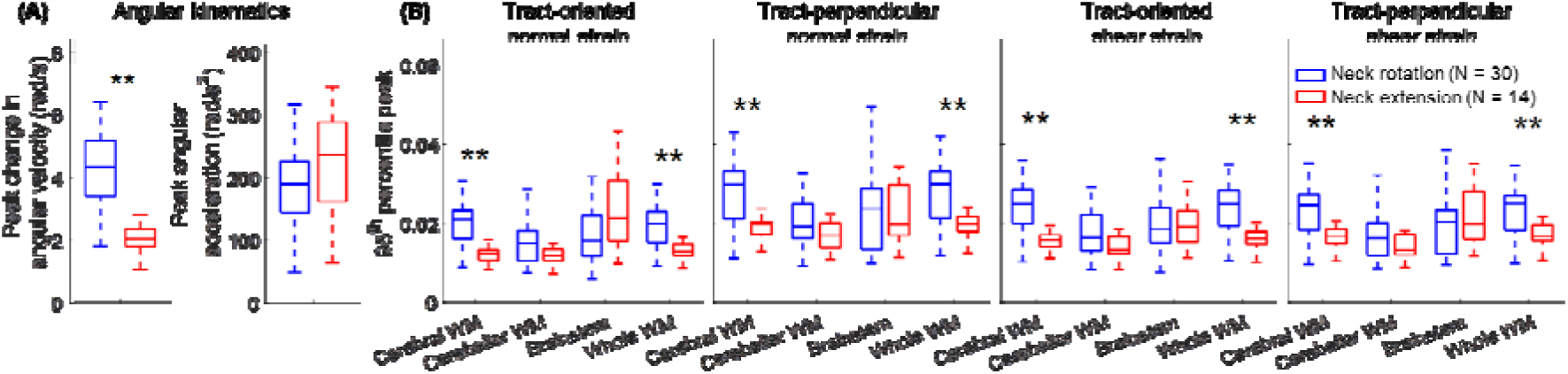
Influence of head loading mode on tract-related strains. (A) Comparisons of peak angular kinematics. (B) Comparison of four tract-related strain peaks (95^th^ percentile values) in cerebral white matter (WM), cerebellar WM, brainstem, and the whole WM between neck rotation and neck extension (\*\**p*<0.001, Mann-Whitney U test).

Pearson’s correction results between angular kinematics peaks (i.e., peak changes in angular velocity and peak angular acceleration) and maximum tract-related strain peaks (95^th^ percentile values) are summarized in Table 1. The correlation results are mainly dependent on the WM region and head loading mode. For the peak change in angular velocity, significant correlations are noted in cerebral WM and whole WM (all strain types) for neck rotation (Table 1A), and in the brainstem (all strain types) for neck extension (Table 1B). When switching the kinematic parameters to peak angular acceleration, similar correlation results are noted, except that the Person’s correlation coefficient values change slightly (Table 1C-D).

**Table 1.** Person’s correlation coefficients between peak angular kinematics (i.e., peak change in angular velocity, peak angular acceleration) and maximum tract-related strains (95^th^ percentile peak) in cerebral white matter (WM), cerebellar WM, brainstem, and whole WM during neck rotation (A, C) and neck extension (B, D). For each loading mode, the significance level is adjusted using the Benjamini-Hochberg method for the correction for multiple comparisons.

|  | Pearson's correlation coefficient values |  |  |  |  |  |  |  | Pearson's correlation coefficient values |  |  |  |  |  |  |  |
| --- | --- | --- | --- | --- | --- | --- | --- | --- | --- | --- | --- | --- | --- | --- | --- | --- |
|  | Tract-related strains vs. peak change in angular velocity |  |  |  |  |  |  |  | Tract-related strains vs. peak angular acceleration |  |  |  |  |  |  |  |
|  | (A) Neck rotation (N=30) |  |  |  | (B) Neck extension (N=14) |  |  |  | (C) Neck rotation (N=30) |  |  |  | (D) Neck extension (N=14) |  |  |  |
| Tract-oriented normal strain | 0.46 | 0.10 | 0.07 | 0.43 | 0.14 | 0.53 | 0.67 | 0.32 | 0.46 | 0.15 | 0.07 | 0.43 | 0.15 | 0.51 | 0.65 | 0.32 |
| Tract-perpendicular normal strain | 0.52 | 0.12 | -0.02 | 0.48 | 0.20 | 0.46 | 0.68 | 0.31 | 0.53 | 0.14 | 0.01 | 0.50 | 0.19 | 0.46 | 0.67 | 0.31 |
| Tract-oriented shear strain | 0.56 | 0.17 | 0.02 | 0.52 | 0.16 | 0.60 | 0.75 | 0.32 | 0.57 | 0.18 | 0.01 | 0.53 | 0.15 | 0.61 | 0.77 | 0.31 |
| Tract-perpendicular shear strain | 0.52 | 0.12 | 0.06 | 0.49 | 0.24 | 0.48 | 0.70 | 0.34 | 0.53 | 0.12 | 0.08 | 0.50 | 0.24 | 0.45 | 0.72 | 0.33 |
|  | Cerebral WM | Cerebellar WM | Brainstem | Whole WM | Cerebral WM | Cerebellar WM | Brainstem | Whole WM | Cerebral WM | Cerebellar WM | Brainstem | Whole WM | Cerebral WM | Cerebellar WM | Brainstem | Whole WM |

## Discussions

Using the data of 42 volunteer impacts from the open-accessed BBIR dataset,^44, 48^ the current study estimates *in vivo* WM fiber deformation based on four tract-related strains by relating the brain strain tensor measured by tMRI to fiber orientation estimated from DTI. This multimodal imaging analyses deliver four strain parameters (i.e., tract-oriented normal strain, tract-perpendicular normal strain, tract-oriented shear strain, and tract-perpendicular shear strain), collectively representing a comprehensive description of the fiber deformation. Our work finds the tract-perpendicular normal strain is the largest among the four strain types, unveiling fundamental new aspects of WM dynamics. The reported strain results can be used to evaluate the biofidelity of computational human head models, especially those intended to predict tract-related strains under non-injurious impacts.

The range of tract-related strains in our *in vivo* analyses are generally agreed with the regime in one early *in vivo* measurement of brain strain, and far smaller than the proposed brain injury thresholds from *in silico* studies and the strain magnitude in *in vitro* studies with axonal pathology (Table 2). In 2020, Knutsen and coworkers^48^ presented the first *in vivo* measurement of tract-oriented normal strain and reported the range to be 0.007-0.029, largely overlapping with the values in our study (0.009-0.030). *In silico* models have been used to develop brain injury thresholds based on the tract-oriented normal strain. For example, by relating binary injury outcomes (i.e., concussion or no concussion) in sports-related impacts with continuous tract-oriented normal strain data from computational simulations, Giordano and Kleiven^60^ proposed an injury threshold of 0.09 for the corpus callosum and 0.15 for the brainstem. Hajiaghamemar and Margulies^29^ co-registered the numerically derived tract-oriented normal strain contour with the histology-identified neuropathological map in piglets, reporting the axonal injury threshold fell within the range of 0.08-0.14. Similar thresholds were also developed for diffuse axonal injury in humans (0.15),^61^ brain injury in nonhuman primates (0.12 for mild trauma and 0.19 for severe trauma),^62^ and impact-induced behavioural changes in rats (0.09).^63^ A few *in vitro* studies linked the strain at the cultured tissue to the injury outcome. For example, LaPlaca and colleagues^64^ found a significant loss of neurite that occurred at the conditions when the tract-oriented shear strain peaked (∼0.56). In another *in vitro* model^65^ with neuronal death, the tract-oriented normal and shear strains were within the range of 0.12-0.64 and 0.11-0.48, respectively. Bar-Kochba and coworkers^66^ found the axonal bleb formation was strongly correlated with the tract-oriented shear strain with a threshold of 0.14. Nakadate and coworkers^67^ found swelling in neurons aligned with and perpendicular to the loading direction (a strain of 0.22 at a strain rate of 37 /s). Braun and coworks^68^ also observed tau pathology in neurons oriented along and perpendicular to the loading direction at the strain magnitudes of 0.05 and 0.20, respectively. It should be clarified that the scale of interest may not necessarily be the same between the *in vitro* studies (axon/neuron level) and the current work (fiber tract level). Taken together, the tract-related strain values in *in silico* and *in vitro* studies with varying injury outcomes largely exceed the range in our study under *in vivo* volunteer impacts.

**Table 2.** Comparison of tract-related strains among *in vivo*, *in silico*, and *in vitro* studies. Note that the nomenclature in the listed studies may not be the same and are converted to the one in the current study. For the three superscripts, ^a^: the values are estimated from Fig 7 of the cited reference; ^b^: the values are estimated from Fig 4 of the cited reference; ^c^: the values are estimated from Fig 9 of the cited reference; ^d^: In the original article, the value was averaged along the length of the neuron; ^e^: the values are obtained based on the assumption that the strain levels in the embedded neuron are the same as those imposed to the cultured experimental tissue.

| Reference |  | Peak angular acceleration<br>(rad/s<sup>2</sup>) |  | Tract-oriented<br>normal strain | Tract-perpendicular<br>normal strain | Tract-oriented<br>shear strain | Tract-perpendicular<br>shear strain |
| --- | --- | --- | --- | --- | --- | --- | --- |
| In vivo | Current study | Neck rotation<br>(N=30) | 185.5 ±<br>59.2 | 0.009-0.030 | 0.011-0.042 | 0.010-0.035 | 0.010-0.035 |
|  |  | Neck extension<br>(N=14) | 228.6 ±<br>8.3 | 0.009-0.017 | 0.012-0.024 | 0.010-0.020 | 0.011-0.022 |
|  | Knutsen and<br>coworkers<sup>48</sup> | Neck rotation<br>(N=10) | 212.8 ±<br>32.1 | 0.007-0.024<sup>a</sup> |  |  |  |
|  |  | Neck extension<br>(N=10) | 282.8 ±<br>47.4 | 0.008-0.029<sup>a</sup> |  |  |  |
| Reference |  | Thresholds of 50% injury<br>risk |  | Tract-oriented<br>normal strain | Tract-perpendicular<br>normal strain | Tract-oriented<br>shear strain | Tract-perpendicular<br>shear strain |
| In silico | Giordano and<br>Kleiven<sup>60</sup> | Concussion (human) |  | 0.07-0.15 |  |  |  |
|  | Sahoo and<br>colleagues<sup>61</sup> | Diffuse axonal injury<br>(human) |  | 0.15 |  |  |  |
|  | Wu and colleagues<sup>62</sup> | Brain injury (primate) |  | 0.12-0.19 |  |  |  |
|  | Hajiaghdamemar and<br>Margulies<sup>29</sup> | Diffuse axonal injury (pig) |  | 0.08-0.14 |  |  |  |
|  | Premi and<br>colleagues<sup>63</sup> | Behavioural alteration (rat) |  | 0.09 |  |  |  |
| Reference |  | Injury pathology |  | Tract-oriented<br>normal strain | Tract-perpendicular<br>normal strain | Tract-oriented<br>shear strain | Tract-perpendicular<br>shear strain |
| In vitro | LaPlaca and<br>coworkers<sup>64</sup> | Neurite loss |  |  |  | ~0.56<sup>b</sup> |  |
|  | Cullen and<br>LaPlaca<sup>65</sup> | Neuron death |  | ~0.12-0.64<sup>c</sup> |  | ~0.11-0.48<sup>c</sup> |  |
|  | Bar-Kochba and<br>coworkers<sup>66</sup> | Neuron bleb formation |  |  |  | 0.14<sup>d</sup> |  |
|  | Nakadate and<br>coworkers<sup>67</sup> | Neuron swelling |  | 0.22<sup>e</sup> | 0.22<sup>e</sup> |  |  |
|  | Braun and<br>coworks<sup>68</sup> | Tau mislocalization |  | 0.05<sup>e</sup> | 0.2<sup>e</sup> |  |  |

Our work unravels the WM fiber tract deforms most in normal in the perpendicular direction, holding significant implications for brain injury mechanics. At present, the stretch along the fiber direction is widely recognized as the pathogenesis of axonal injury, as substantiated by the growing body of literature that employs tract-oriented normal strain as an axonal injury criterion.^47, 60–62, 69, 70^ Indeed, several *in vitro* studies reported under the same magnitude level, the tract-oriented normal strain instigated more severe axonal pathology than the tract-perpendicular normal strain,^67, 68^ indicating a higher tolerance level for the normal strain perpendicular to the neurites. However, our work finds that under *in vivo* human head impacts, the tract-perpendicular normal strain is significantly larger than the tract-oriented normal strain. Thereby, one cannot exclude the possibility that the tract-perpendicular strain is one possible instigator of axonal pathology in real-world impacts. When switching to the shear strain, *in vitro* studies also observed neurite loss and neuron death in the regime when the tract-oriented shear strain peaked.^64, 65^ Taken together, it can be concluded that the existing brain injury investigation may have neglected important aspects of axonal pathogenesis.

The larger value of tract-perpendicular strain than its counterparts under volunteer tests may be attributed to the anisotropic material property of WM. Of particular relevance, several *in vivo* studies on the human^71, 72^ and the minipig^73, 74^ analyzed the wave motion fields measured by MRE using transversely isotropic nonlinear inversion algorithms and found that the tensile stiffness perpendicular to the fiber tract was lower than that parallel to the fiber tract. Similar findings have also been reported in several *in vitro* brain tissue material tests.^75–79^ The mechanical reinforcement along the fiber direction provides a possible explanation for why the fiber tract deforms more in the perpendicular direction than the normal direction. However, it should be noted that no consensus has been reached yet about brain anisotropy as another body of literature reported that no significant mechanical dependency on WM fiber orientation was noted in the brain tissue.^80–83^ Another potential explanation for disproportional magnitudes of tract-related strains is related to the feature of strain tensor. For the tract-oriented normal strain, its magnitude is constant across the tract-aligned coordinate system with one axis aligned with the fiber direction. In contrast, the magnitude of the tract-perpendicular strain changes when the coordinate system rotates around the fiber direction and its maximum value across all tract-aligned coordinate systems is presented in the current study, corresponding to the most severe loading scenario (Appendix A).

Our results also find laterally symmetric strain patterns with respect to the midsagittal plane under neck extensions, but not under neck rotations (Figs 3-4). This can be explained by the presence of cerebral falx at the interhemispheric fissure with mechanically stiffer properties than the brain (i.e., ∼ 22.9 MPa for the elastic moduli of falx^84^ vs. ∼ 1 kPa level for the shear stiffness of the brain).^85^ During neck extensions, the brain primarily moves within the sagittal plane (confirmed by the displacement field recorded by tMRI) with little interaction with the anterior-posterior oriented falx, and hence left-right symmetric deformation patterns are developed in the brain. Under neck rotations, the primary brain displacement is noted within the coronal plane. The presence of the mechanically stiffer falx disturbs the brain motion, leading to asymmetric strain patterns. Similar findings have been reported in earlier *in vivo* imaging studies^38, 48^ and *in silico* impact simulations.^86, 87^ Collectively, these studies highlight the mechanical role of the falx during impacts.

Our study represents a significant extension of one early *in vivo* tract-oriented normal strain estimation (10 neck rotations and 10 neck extensions)^48^ by involving more impacts (30 neck rotations and 14 neck extensions). Except for the overlap of the tract-oriented normal strain ranges described above, both our work and the early study^48^ found high tract-oriented normal strains primarily in the cerebral WM for neck rotations and in the brainstem for neck extensions (Fig 3), providing independent verification of each other. Moreover, we generate spatially detailed information on how the axonal fiber responds to normal strain perpendicular to the fiber tract and shear strain along and perpendicular to the fiber tract, representing three new aspects of fiber deformation. Knutsen and coworkers^48^ found the square of angular velocity peak strongly correlated with the maximum tract-oriented normal strain in the corpus callosum during neck rotations and in the brainstem during neck extensions. Here, we expand the correlation by incorporating other kinematic-based metrics (i.e., peak change in angular velocity and peak angular acceleration) and four tract-related strains, through which we find both angular kinematics-based peaks significantly correlated with the tract-related strains (all four types) in cerebral WM and whole WM for neck rotations, and in the brainstem (all four types) for neck extensions (Table 1). To provide further verification of the observed relationships, in-house *in silico* work is ongoing to extend the correlation analyses by including impacts with higher severities.^88^

The current work provides important data for the validation of computational head models. Partially owing to the development of numerical algorithms (e.g., finite element method and material point method), computational head models have become increasingly used to study brain biomechanics.^16^ Prior to the utilization, the reliability of computational head models requires to be extensively evaluated against relevant experimental measurements. Existing head models are often validated against cadaveric measurements in the form of brain-skull relative motion^7, 11, 89^, maximum principal and shear strain,^8, 9^ and maximum principal strain rate^10^ from only a handful of intracranial sites with the impact severities close to traumatic levels. Despite the great value of cadaveric tests, the material property of the brain changes after death. This is further complicated by the influence of post-mortem time and storage conditions without consensuses being reached yet.^85^ One possible solution is to validate the model’s responses collectively against the measurements from high-severity, sparsely recorded cadaveric experiments and low-severity, spatially detailed volunteer tests (such as the brain strain data presented herein). This may be particularly relevant for those models with intended usage to predict the brain response under non-injurious impacts. Emerging attempts have been noted in validating the computational head model against *in vivo* brain responses in the form of brain-skull relative displacement,^90^ maximum principal strain,^22, 23, 87, 91^ maximum shear strain,^87^ and tract-oriented normal strain.^22, 23^ Our study computes 3D normal and shear strains along and perpendicular to the fiber tract (spatial resolution: 1.5 mm or 2 mm, temporal resolution: 18 ms, 19.5 ms, or 20 ms) across the whole WM from 44 volunteer impacts. These strain data are complementary to the existing cadaveric experiments. Collectively, these *in vivo* and *in situ* experimental data may contribute to the validation of computational brain models towards a comprehensive degree and ensure the confidence of model usage to estimate brain response across the sub-injurious and injurious levels.

As motivated by the prevalence choice of using normal and shear strain (e.g., maximum principal strain, maximum shear strain) for brain injury analysis,^92^ the current study presents the normal strain and shear strain along and perpendicular to fiber tracts. Recent trends were noted in employing the compressive normal strain as the parameter of interest, e.g., minimum principal strain,^93–95^ compressive strain at the fiber tract level,^96^ neuronal level,^66^ and microtube level.^97^ For example, two *in vitro* studies imposed compressive normal strains perpendicular to the main axonal direction and found cytoskeletal degeneration (e.g., changes in neurofilament and microtube distributions),^98, 99^ suggesting the tract-perpendicular normal strain in compression as another possible trigger of axonal injury. In relation to our study, the tract-perpendicular normal strain at a given time instance may attain both positive value (i.e., tensile strain) and negative value (i.e., compressive strain) with dependency on the choice of coordinate systems (see equations 3-4 in Appendix A). Although the current study focuses on the tensile normal strain (e.g., *_ε_*^max^ represents the maximum tract-perpendicular normal strain, see Appendix A), we also present the equation for computing the minimum tract-perpendicular strain across all tract-aligned coordinate systems (see *ε*^min^ in Appendix A). Other researchers, who are interested in computing the compressive normal strain perpendicular to the fiber tract, may find this equation helpful.

The current study is based on volunteer tests and imaging data from the open-access BBIR dataset. Despite its great value, the dataset suffers certain limitations (e.g, varying resolutions in the temporal and spatial domains between different tests) that were already detailed by the dataset developer.^44^ Other challenges and caveats, e.g., the limited number of image frames in tMRI,^41, 48^ the adverse effect of decreased signal-to-noise ratio on strain estimation as tag lines fade,^100^ incapability of diffusion tensor to describe the region of crossing fibers with multiple orientations present within the voxel,^48, 101, 102^ are exhaustively discussed elsewhere and not reiterated herein.

## Conclusion

Based on the volunteer impact data from the BBIR dataset,^44, 48^ this study calculates WM tract-related deformation during voluntary impacts by relating the tMRI-recorded brain strain tensor to DTI-derived fiber orientation through coordinate transformation. By extending the previous effort that focused on the normal strain along the fiber tract, we present *in vivo* data of the normal strain perpendicular to the fiber tract and shear strain along and perpendicular to the fiber tract towards a multifaceted understanding of WM dynamics. Our work finds the fiber tract deforms most in the perpendicular direction, deciphering new fundamentals of brain biomechanics. The reported strain results under volunteer impacts can be used to evaluate the fidelity of computational head models, especially those intended to predict fiber deformation under non-injurious conditions.

## Transparency, Rigor, and Reproducibility

The *in vivo* imaging data used for the computation of white matter fiber deformation are obtained from an open-access dataset (see the acknowledgements). The rigor and reproducibility are ensured by the steps for the calculation of four tract-related strains and statistical analyses presented in the paper. The yielded strain maps, including 4 tract-related strains from 44 impacts, can be requested by emailing request to the corresponding author.

## Acknowledgements

Data were provided in part by the Brain Biomechanics Imaging Repository (http://www.nitrc.org/projects/bbir). Principal Investigators: P. Bayly, D. Pham, C. Johnson, J. Prince, K. Ramesh, through grants U01 NS11212 and R56 NS055951. The content of this article is solely the responsibility of the authors and does not necessarily represent the official views of neither funding agencies nor the dataset developer. The computation of tract-related strains was enabled by resources provided by the National Academic Infrastructure for Supercomputing in Sweden (NAISS) at the center for High Performance Computing (PDC) partially funded by the Swedish Research Council through grant agreement no. 2022-06725.

## Author contribution

**Zhou Zhou**: Conceptualization, Formal analysis, Funding acquisition, Methodology, Software, Visualization, Original draft, Review & editing. **Christoffer Olsson**: Conception and study design, Investigation, Software, Review & editing. **T. Christian Gasser**: Formal analysis, Investigation, Methodology, Original draft, Review & editing. **Xiaogai Li**: Visualization, Methodology, Review & editing. **Svein Kleiven**: Methodology, Original draft, Review & editing, Funding acquisition.

## Appendix A Computation of tract-related strains

At a given time instance, the white matter (WM) tract-related strains are calculated by transforming the brain strain tensor to the coordinate system with one axis aligned with the fiber orientation. For a WM voxel (i.e., voxel with the fractional anisotropy value ≥0.2), the primary fiber orientation is approximated as the first principal eigenvector of diffusion tensor (denoted as ***v*** = _[_*v*_1_ *v*_2_ *v*_3_]^T^). At each time frame, the brain strain tensor ***^E^***_global_ in tMRI (Fig 1D) is expressed with reference to the global coordinate system (***XYZ***, ***X*** = [1 0 0]^T^, ***Y*** =[0 1 0]^T^, ***Z*** =[0 0 1]^T^, Fig A1A). We rotate ***E***_global_ into the local coordinate system (1) (***xyz*** in Fig A1B, ***x*** is aligned with ***v***, ***y*** is the cross product of ***x*** and ***X***, ***z*** is the cross product of ***x*** and ***y***) via ***E*** _track_ = ***QE***_global_ ***Q***^T^, in which

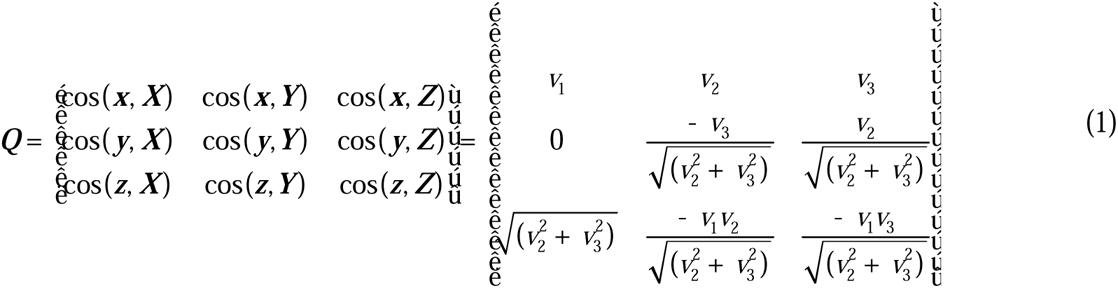

The six independent strain components in the local coordinate system (1) are illustrated in Fig A1C.

With respect to strain components *ε_xx_*, *γ*_*xy*_, and *γ_xz_* (the latter two act at the plane perpendicular to the fiber direction, i.e., *yz* plane, see Fig A1D), we form two strain measures, i.e., the tract-oriented normal strain *ε_a_* = *ε_xx_* represents the normal strain along the fiber tract (Fig A1E), and the shear strain 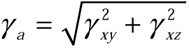 within the *yz* plane. Given the symmetry of the strain tensor, *γ_a_* also appears along the fiber and it therefore denotes the tract-oriented shear strain (Fig A1E).

Now we consider the strain components _*ε_yy_*_, *ε_zz_*, and *γ_yz_*, acting on the *xy* and *xZ* planes (Fig A1C). Rotating these planes around the fiber direction changes the values of the individual strain components (i.e., 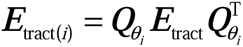 in the local coordinate system (*i*) (***xy****_i_****z****_i_*)) with dependency on the rotating angle (0 ^°^ ≤ *θ*_*i*_ ≤ 360^°^) (see Fig 5B). Note that,

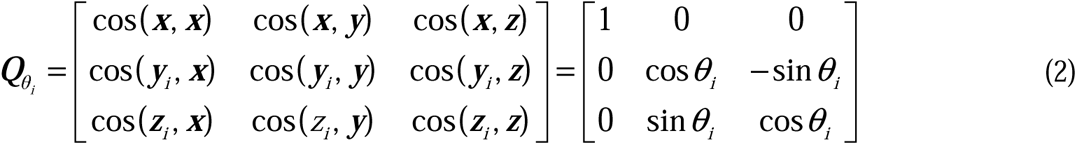

The equations for the strain components *ε*_*yy*_, *ε_zz_*, and *_γ yz_* in ***E***_tract(*i*)_

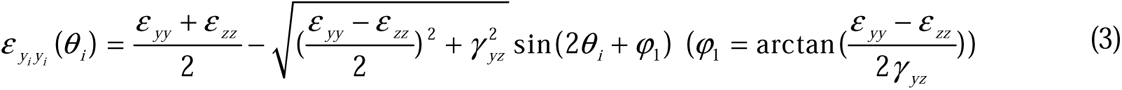

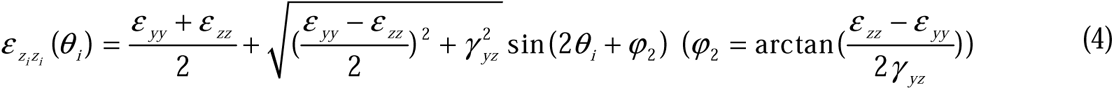

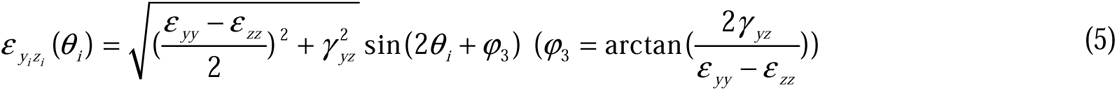

Based on equations 3-4, 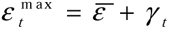 denotes the maximum tract-perpendicular normal strain and 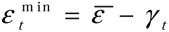 denotes the minimum tract-perpendicular normal strain, where 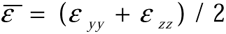. Note that in the main text, the 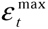 (i.e., the maximum tract-perpendicular normal strain) are reported.

Based on equation 5, we determine the maximum tract-perpendicular shear strain 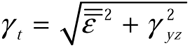, where 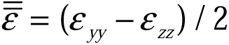.

**Fig A1.**
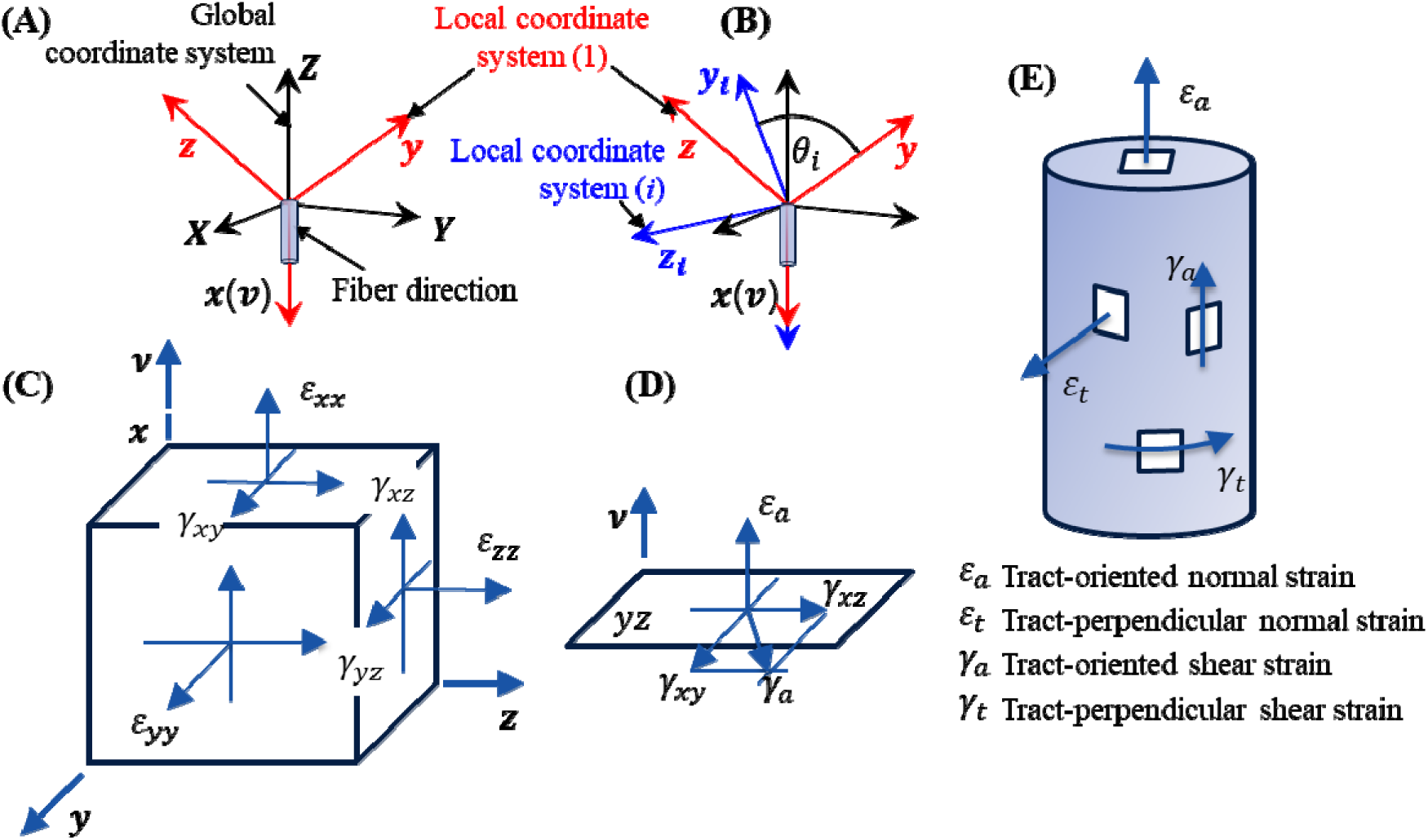
Computation of four tract-related strains via coordinate transformation. (A) Rotate the strain tensor from the global coordinate system (***XYZ***) to the local coordinate system (1) (***xyz***), in which ***x*** represents the fiber direction (***v***). (B) Rotate the strain tensor from the local coordinate system (1) to the local coordinate system (*i*) (***xy****_i_* ***z****_i_*). Note that the local coordinate system (*i*) is obtained by rotating the local coordinate system (1) along the fiber direction (***V***) by arbitrary degree (*θ_i_*). (C) The six independent strain components in the local coordinate system (1). (D) Illustration of *ε_xx_*, *γ_xy_*, and *γ_xz_*, of which the latter two act in the *yz* plane. (E) Illustration of four tract-related strains that are parameters of interest in the current study, i.e., tract-oriented normal strain *ε_a_* characterizes the normal strain along the fiber tract, tract-perpendicular normal strain *ε_t_* characterizes the maximum normal strain perpendicular to the fiber tract, tract-oriented shear strain *γ_a_* characterizes the shear strain along the fiber tract, and tract-perpendicular shear strain *γ_t_* characterizes the maximum shear strain perpendicular to the fiber tract.

The presented method has been similarly used to compute the orientation-dependent strains in neural cells in *in vitro* models.^64–66, 103^ We previously used this method to estimate the tract-related deformation during *in silico* impact simulations, in which the real-time fiber orientation (i.e., the deformed fiber orientation that changes at each timestep of the simulation) was used.^51, 52^ We acknowledge that is flawed and the undeformed fiber orientation should be used (e.g., the static first principal eigenvector of diffusion tensor used in the current study). Errata to our previous studies^51, 52^ will be published separately.

## Appendix B Influence of the disparity of voxel resolution on white matter tract-related strains

To assess how the difference in voxel resolutions (1.5 mm vs. 2.0 mm) affects the tract-related strains, strain tensor data from one randomly selected volunteer test with a spatial resolution of 1.5 mm is down-sampled to 2 mm. Both the original and down-sampled imaging data are used to calculate the tract-related strains (Fig B1). The similarity in strain contours and overlap in the percentile strain values between two resolutions indicate the spatial resolution disparity (i.e., 1.5 mm vs. 2.0 mm) has negligible effects on four maximum tract-related strains.

**Fig B1.**
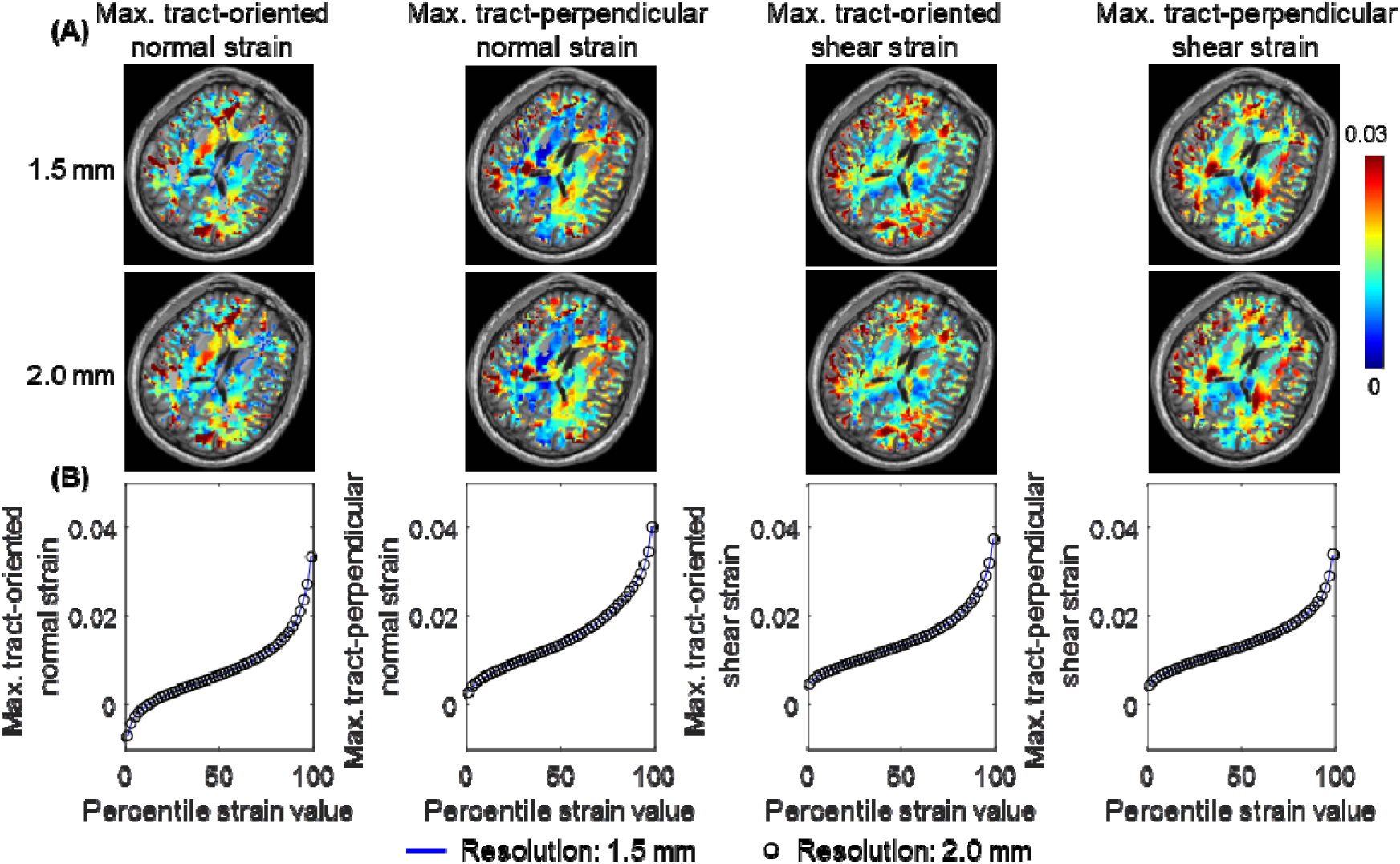
Influence of imaging down-sampling (from 1.5 mm to 2.0 mm) on the maximum values of four tract-related strains in terms of strain distribution (A) and percentile strain values (B).

## Appendix C Comparisons of four tract-related strain peaks

**Fig C1.**
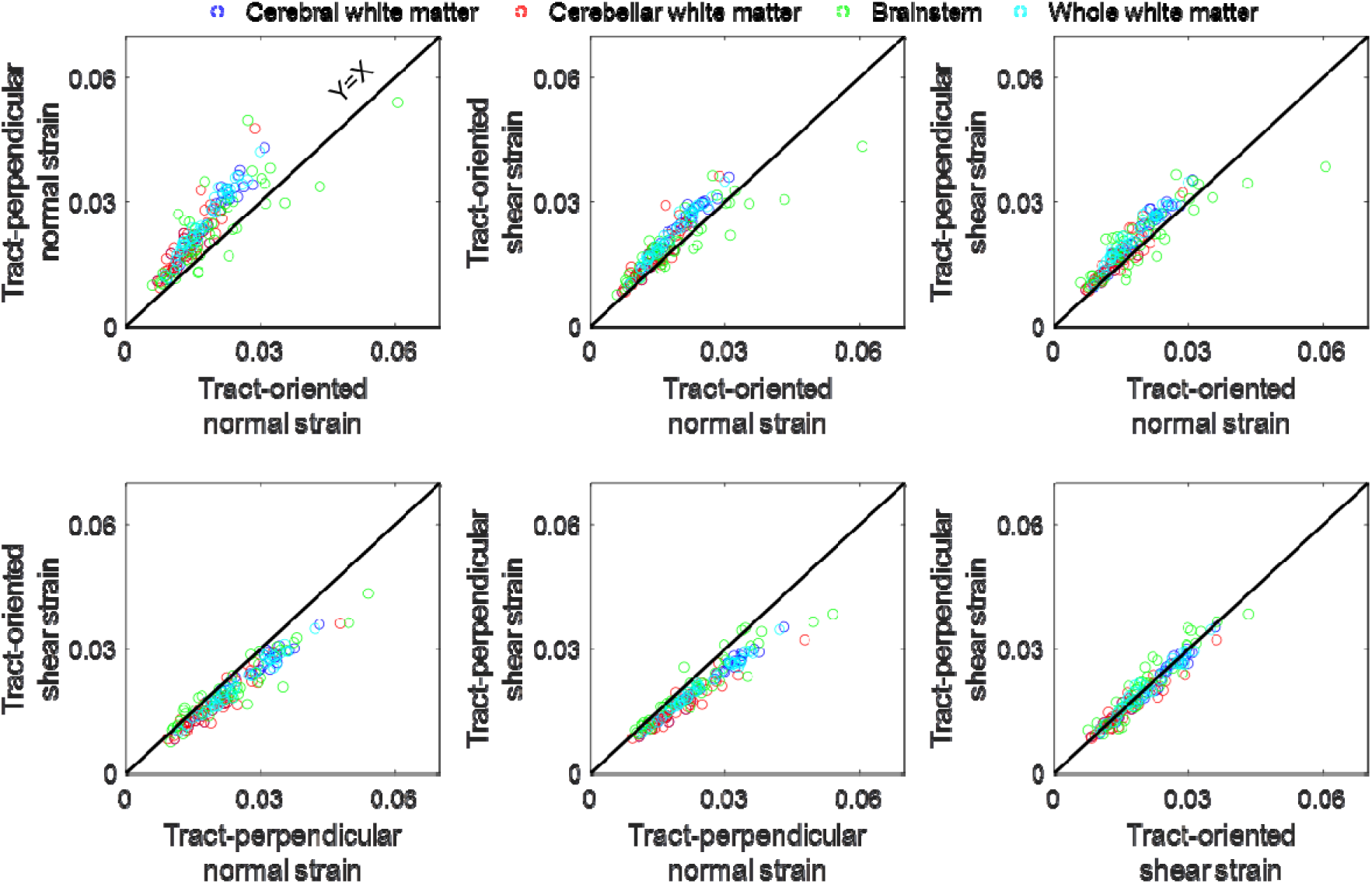
Scatter plots of the peaks of four tract-related strains in the cerebral white matter, cerebellar white matter, brainstem, and whole white matter.

